# *KaMRaT*: a C++ toolkit for *k*-mer count matrix dimension reduction

**DOI:** 10.1101/2024.01.15.575511

**Authors:** Haoliang Xue, Mélina Gallopin, Camille Marchet, Ha N. Nguyen, Yunfeng Wang, Antoine Lainé, Chloé Bessiere, Daniel Gautheret

## Abstract

**Summary:** *KaMRaT* is a program for processing large *k*-mer count tables extracted from high throughput sequencing data. Major functions include scoring *k*-mers based on count statistics, merging overlapping *k*-mers into longer contigs and selecting *k*-mers based on their presence in certain samples. *KaMRaT* ‘s main application is the reference-free analysis of multi-sample and multi-condition datasets from RNA-seq, as well as ChiP-seq or ribo-seq experiments. *KaMRaT* enables the identification of condition-specific or differential sequences, irrespective of any gene or transcript annotation.

**Implementation and availability:** *KaMRaT* is implemented in C++. Source code and documentation are available via https://github.com/Transipedia/KaMRaT. Container images are available via https://hub.docker.com/r/xuehl/kamrat.

## Introduction

RNA-seq data analysis commonly involves comparison of sequence reads to a reference genome or transcriptome and quantification of annotated genes or transcripts (Van den Berge et al., 2019). While convenient, this approach ignores a wide range of variations present in the original sequence data. These variations may come from novel RNA isoforms, RNAs from repeats, intergenic regions and exogeneous species such as viruses, as well as from RNAs with small variations such as SNPs and indels. An emerging strategy to investigate all possible RNA variations at once is to use *k*-mers. First a *k*-mer counter, e.g. (Marçais and Kingsford, 2011; Lemane et al., 2022b), extracts and counts all successive substrings of length *k* from the raw sequence reads. Various pipelines use *k*-mer count tables to select biologically relevant *k*-mers, possibly combining *k*-mers into longer contigs (Audoux et al., 2017; Rahman et al., 2018; Lorenzi et al., 2020; Lemane et al., 2022a). However, pipelines are cumbersome and slow to run. We felt that a standalone program performing common operations on *k*-mer count tables, such as *k*-mer selection and contig assembly would facilitate a greater diffusion of *k*-mer analysis among RNA-seq users. We thus developed *KaMRaT* (*k*-mer Matrix Reduction Toolkit), a lightweight and multi-functional toolkit implemented in C++ for *k*-mer matrix analysis, filtering and contig assembly.

## Program Description

*KaMRaT* takes as input a *k*-mer count table generated by any *k*-mer counter with *k*-mers as the first column. Other types of features are allowed (for instance gene IDs, sequence contigs) for certain operations, as long as they are also provided as the first column of the table. The program is composed of 6 main modules (Figure 1 and suppl. Methods).

**Figure 1:**
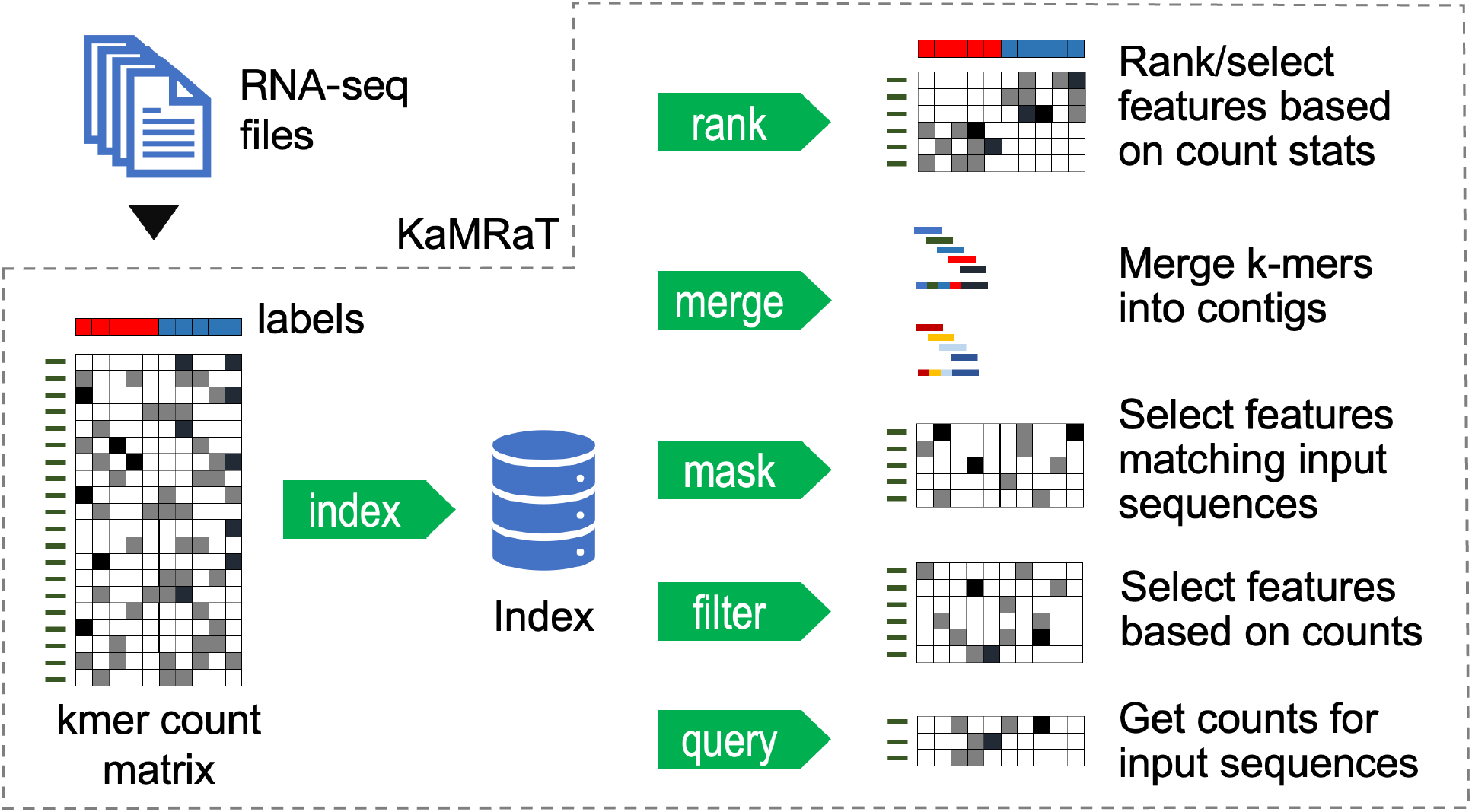
*KaMRaT* modules. Once an index is created, modules can be combined in different order, such as *score*+*merge*. Modules *mask* and *query* require a set of sequences as further input. Module *merge* can also output a new count table for the contigs.

- *index* creates a binary index of the features and counts on disk. This allows subsequent modules to randomly access count vectors without parsing the whole table.
- *score* scores and selects features based on univariate statistical tests, using categorical or numerical column labels provided as input. Available tests include T-test, signal-to-noise ratio (Golub et al., 1999), Detection of Imbalanced Differential Signal (DIDS) (de Ronde et al., 2013), logistic regression accuracy, Bayes classifier accuracy, Pearson and Spearman correlation. Label-free tests are also available, including absolute and relative standard deviation and information entropy.
- *merge* merges *k*-mers into contigs. By default, overlapping *k*-mers are iteratively merged until an ambiguity is met or no more *k*-mer satisfying a given minimum overlap length is available, similar to “unitigs” (Pevzner et al., 2004). An optional “intervention” mode uses correlation between count vectors to determine whether two *k*-mers or contigs with an acceptable overlap should be merged.
- *filter* deletes or selects features based on counts and recurrence thresholds.
- *mask* deletes or selects features matching given fasta sequences.
- *query* estimates count vectors of an input list of sequences based on their constitutive *k*-mers.

## Performance overview

We assessed *KaMRaT* on simulated and real datasets of up to 150 RNA-seq samples. The index size is about 300Gb for 150 samples (Fig. S1A), i.e. equivalent to 1/3 of the original compressed fastq files. Once the index is created, *score* operations run in linear time and memory, allowing to process 420M *k*-mers or contigs in 85min on a single processor with only 6Gb RAM. The most ressource-intensive module, *merge* runs in log-linear time and can process 420M *k*-mers in about 3h, using 80Gb of RAM (Fig. S1B). Note that large *k*-mer tables can be first processed with *score* so that only a subset of high scoring *k*-mers are fed to *merge*, thus considerably reducing the memory requirement.

A key contribution of *KaMRaT* is the intervention mode enabling a significant reduction of misassemblies when merging *k*-mers. Using *k*-mers from simulated reads or real RNA-seq reads from human tissues followed by T-test selection (supplementary material), the various intervention options changed 5% to 25% of the output contigs compared to intervention-free *k*-mer extension (Fig. S2A). Contigs produced using any of the intervention options were shorter but had higher rates of perfect alignment when aligned to reference transcripts (Fig. S2B-C).

## Applications

Typical *KaMRaT* applications are briefly presented below. Supplementary material and the *KaMRaT* github repository https://github.com/Transipedia/KaMRaT/tree/master/toyroom include example workflows and a toy dataset.

### Supervised feature selection

This is the first intended application of *KaMRaT*, typically achieved using an *index-score-merge* pipeline. *K*-mers of interest (for instance differentially expressed) are selected using any of the supervised test provided in the *score* module, and selected *k*-mers are merged into contigs.

### Unsupervised feature selection

*KaMRaT score* supports unsupervised feature selection using standard deviation and information entropy. These methods can help reduce a *k*-mer table dimension independently of the variable to be predicted, thus avoiding information leakage in machine learning applications.

### Finding features correlated to another feature

*KaMRaT score* enables retrieving features that correlate with a quantitative target vector, such as time, a measure of drug effect or the expression of a given gene.

### Retrieving condition-specific *k*-mers or contigs

*KaMRaT filter* can be used to identify features expressed exclusively in samples from one condition. For instance, in a dataset with normal and tumor samples a *KaMRaT index-filter-merge* workflow can retrieve tumor-exclusive contigs.

## Perspectives

*KaMRaT* offers a unique suite of tools for studying feature dimensionality and RNA variations. Three modules are pivotal: *score, merge* and *filter. KaMRaT score* integrates currently 12 methods to reduce feature dimensionality in a supervised or unsupervised fashion, which should fit multiple research situations. *KaMRaT merge* builds on the concept of local *k*-mer extension (“unitigs”) to improve extension precision by leveraging count data. By our tests, intervention significantly improved extension correctness. However, these contigs do not compare to those produced by a full-length transcript assembler in that they are interrupted whenever ambiguities occur in the graph, for instance when encountering a SNP. *KaMRaT filter* allows retrieval of condition-specific features, which can be useful for collecting all RNA variations that are specific to a given sample set.

Although designed primarily for *k*-mer matrices, the *score* and *filter* modules apply to any generic count matrix such as gene-/transcript-expression matrices. This enables building classifiers from reference-free features (*k*-mers, contigs) and reference-based features (genes, transcripts) in a consistent and comparable way.

## Acknowledgements

This work was supported by grants ANR-18-CE45-0020 and ANR-22-CE45-0007. We are thankful to Rayan Chikhi for fruitful discussions and Teo Lemane for help with software engineering.

## Supplementary material for

### Supplementary Methods

#### Workflow overview

*KaMRaT* ‘s workflow can be described as follows: several samples (definition 1) are processed so that each feature (definition 4) is associated to a sample count vector (definition 5). The resulting feature count matrix (definition 6) is the input of the *KaMRaT* workflow. An index is then computed over the matrix, which can be processed through different operations. Thereby, the initial matrix is reduced into a sub-matrix where features are more relevant to a research goal, and - if features are *k*-mers (definition 2) - less interdependent. In a nutshell, given a dataset of which the count matrix has dimension *P* × *N, KaMRaT* aims to reduce the matrix to dimension *p* × *N* where *p < P*.

#### Definitions

##### Definition 1: Sample and dataset

*A sample may correspond to one (in single-end sequencing) or two (in paired-end sequencing) FASTQ files. Multiple samples used in the same analysis compose a dataset. Let N denote the number of samples in the dataset for KaMRaT processing. Samples are numbered from* 0 *to* (*N* − 1).

##### Definition 2: *k*-mer and *k*-mer count

*k-mers are successive, overlapping substrings of a fixed length k extracted from sequencing reads. k-mer occurrences can be counted across samples, which are denoted by k-mer counts hereafter*.

##### Definition 3: *k*-mer merging/extension and *k*-mer contigs

*Merging and extension are two equivalent terms, describing the process of aggregating overlapping k-mers to produce a single, longer sequence, called a k-mer contig, or contig for short*.

##### Definition 4: Feature

*A feature is the abstraction of one property in a dataset. According to the context, a feature can denote a k-mer, a contig, a gene, or a transcript. Let P denote the number of features in the considered dataset*.

##### Definition 5: Sample count vector

*A sample count vector is a numeric vector of size N, which stores a vector of feature counts across samples* {0, …, *N* − 1}.

##### Definition 6: Count matrix

*A count matrix results from stacking sample count vectors of multiple features, where the rows and columns represent features and samples, respectively*.

#### Indexation

To avoid repetitive parsing of the whole matrix by different modules, *KaMRaT* indexes the count matrix that is used for every further operation into binary index files, allowing random access to sample count vectors. Feature names/strings and count vectors are indexed into a single file, with the index positions being stored separately. Downstream modules then only rebuild in memory the association between a feature and the indexed position to its sample count vector, avoiding repetitive processing of count vectors at each subsequent step. Optionally, users may normalize counts at this step based on total counts per sample, following equation (1):

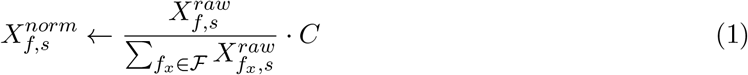

where 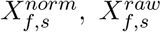: normalized and raw count of feature *f* and sample *s*; ℱ: universe of all features; *C*: constant scaling factor provided by user.

##### Indexation algorithm

*KaMRaT index* generates three files: (i) idx-meta.bin that stores meta-information, (ii) idx-mat.bin which encodes the count matrix, and (iii) idx-pos.bin comprising position for each row of the index matrix in idx-mat.bin, feature by feature. The meta-information includes: sample number, *k*-mer size, *k*-mer strandedness, header row of the input matrix and computed normalization factors. In the case of non-*k*-mer features, the *k*-mer size is set at 0 as a placeholder and strandedness information is ignored. The idx-mat.bin file encodes each feature with its count vector as a row. Each count vector is normalized (if required by user), converted to a character string by bit with *reinterpret cast*, and written into the file followed by the corresponding feature’s sequence/name. The idx-pos.bin file can be organized in two ways according to the value of *k*-mer size. If it equals to 0 (i.e., non-*k*-mer features), it contains a series of position value encoded into character by bit. Otherwise, if features are *k*-mers, it’s composed by a series of pairs encoded from compressed *k*-mer integer code and the corresponding position. The compressed *k*-mer integer code is computed by mapping each nucleotide into two bits: *A* = 00_2_, *C* = 01_2_, *G* = 10_2_, *T* = 11_2_.

##### Note on *k*-mer size

The *k*-mer size is set by the *k*-mer counter and specified at the time of indexation. *KaMRaT* requires that *k*-mers be no longer than 32nt which is the maximum length that an int64 variable can store. There is no lower limit, however for specificity considerations, we recommand to use at least 15nt. Also short *k*-mers would inevitably lead to a higher rate of misassembly by *KaMRaT merge*. Finally, it is recommended to use an odd size number, to avoid potential confusions between a *k*-mer with its reverse-complement when *k* is even (e.g., AAATTT when *k* = 6).

#### Operations

Relying on the binary index of the matrix, *KaMRaT* can perform five operations.

- *score* scores features by their count vectors’ dispersion, informativeness, or association to a given categorical/numerical vector;
- *merge* extends *k*-mers into longer sequences (contigs) based on sequence overlap;
- *filter* extracts/eliminates features according to their counts;
- *mask* retains/removes *k*-mers matching an input sequence list;
- *query* estimates count vectors of a given input list of sequences (*k*-mers or contigs) from the indexed count matrix.

***KaMRaT score*** applies univariate feature scoring by different methods, with an option to select top figures for dimensionality reduction. It scores features either by estimating their count vector’s association to a target vector of the same dimension, or by computing their count vector’s dispersion or amount of information without involving any supplementary variable. The operation allows to select the top *x* or top *x*% features according to user’s indication.

When a target vector is required for association estimation, it is provided through a list of (*sample, value*) pairs in a tabular input file (design file), with the *value* being either categorical (such as samples’ biological conditions) or numerical (such as samples’ estimated immune cell abundance).

Table 1 summarizes available scoring methods with their acceptable target vector type. Detailed information on scoring methods are provided below in this document (*Details on scoring methods*).

**Table 1:** Scoring methods in *KaMRaT score*.

| scorer | target vector type | returned value |
| --- | --- | --- |
| **ttest.padj** | categorical, 2 classes | t-test adjusted p-value with B-H procedure |
| **ttest.pi** | categorical, 2 classes | t-test π-value Xiao et al. (2014) |
| **snr** | categorical, 2 classes | signal-to-noise ratio Golub et al. (1999) |
| **dids** | categorical, $\geq 2$ classes | DIDS de Ronde et al. (2013), adapted |
| **lr** | categorical, 2 classes | logit regression’s accuracy |
| **bayes** | categorical, $\geq 2$ classes | Bayes classifier’s accuracy |
| **pearson** | numerical | Pearson correlation |
| **spearman** | numerical | Spearman correlation |
| **sd** | - | standard deviation |
| **rsd1** | - | relative standard deviation to mean count |
| **rsd2** | - | relative standard deviation to minimum count |
| **entropy** | - | information entropy |

***KaMRaT merge*** performs *k*-mer extension into *contigs* (see definition 3). *k*-mer contigs are obtained using a greedy extension procedure as in (Audoux et al., 2017), i.e., overlapping *k*-mers are iteratively merged until an ambiguity is met (multiple predecessors/successors) or no more *k*-mer satisfying a given minimum overlap length (by default |*k/*2|nt) is available. This extension procedure is conservative : all bifurcations including those due to frequent mutations (such as SNV, indel, etc.) are considered as stop signs. This makes merging quite different from assembly methods such as *rnaSPAdes* (Bushmanova et al., 2019). In principle, this procedure is subject to extension errors, where *k*-mers from independent loci are merged due to their accidental overlap (which frequently occurs in repeats). This led us to propose a novel constraint for local extension: *disagreement of count vectors* (see definition 7).

##### Definition 7: Disagreement of count vectors

*The disagreement of two count vectors is a distance measure used to estimate how likely two count vectors are to be produced by different RNA transcripts*.

When using the *intervention* option of *KaMRaT-merge*, disagreement between count vectors is computed to determine whether two *k*-mers or contigs with an acceptable overlap should be merged. Disagreement is estimated by either Pearson distance, Spearman distance or mean absolute contrast (MAC), defined by equation set (2) where all measures are scaled between 0 and 1. By default, extension is executed when *d*_Pearson_ *<* 0.20.

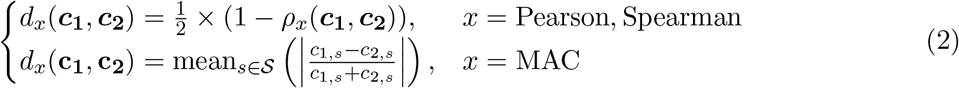

where *d*_*x*_: Pearson/Spearman/MAC distance; **c**_**1**_ and **c**_**2**_: sample count vectors of the two *k*-mers adjacent to the overlapping part, with *c*_1,*s*_ and *c*_2,*s*_ being the components of sample *s*; S: universe of all samples; *ρ*_*x*_: Pearson/Spearman correlation coefficient.

##### Definition 8: Stable merging

*A stable merging is achieved if the resulting contig is determined only by the information at either adjacent side of the overlapping part, independent from information on any other location*.

Stable merging guarantees obtaining the same results regardless of input order of *k*-mers. The original *k*-mer merging procedure, by only considering prefix-suffix overlap, achieves this by nature; however, with sample-count intervention, this does not hold true when count vectors of *k*-mers not located at the merging site are considered for disagreement evaluation. Therefore, to guarantee *stable merging*, disagreement is evaluated only between the two *k*-mers at the merging site.

After a merging operation, the count vector of the resulting contig can be computed for each sample as the mean or median counts of all constituent *k*-mers, according to user’s preference.

***KaMRaT filter*** filters features according to their abundance and recurrence in one or more samples. It can be used to select features expressed above a certain level in one or several samples. Users must provide a design file in which samples are labelled as either “UP” or “DOWN”, and *KaMRaT filter* will select features such that :

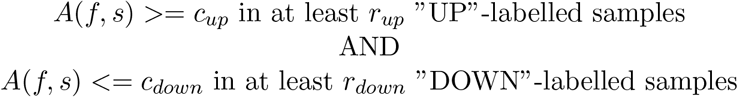

where *A*(*f, s*) is the abundance of a feature *f* in sample *s*, and *c*_*up*_, *c*_*down*_, *r*_*up*_, *r*_*down*_ are provided by users.

A simple use case is the retrieval of features present exclusively in the case condition. Here, control samples are labelled as “DOWN”, with *c*_*down*_ = 0 and *r*_*down*_ = #{control samples}.

*KaMRaT filter* also supports the reverse operation, i.e., to remove the features satisfying given criteria rather than to select them.

***KaMRaT mask*** retains or removes *k*-mers matching an input list of sequences. *k*-mers in the indexed matrix are retained or removed only if they have exact matches in the input sequence.

***KaMRaT query*** estimates count vectors of an input list of sequences based on their constituent *k*-mer counts in the indexed matrix. The query operation has two modes : mean and median, which compute respectively the mean or median count vector of input sequences’ constituent *k*-mers found in the index. If a sequence has no constituent *k*-mer found, it can be either omitted or returned with an all-zero vector, as per user’s preference.

#### Workflow examples

*KaMRaT* ‘s modular design allow users to quickly test and practice different workflows (Figure 1, main text). To produce the input *k*-mer count matrix from individual RNA-seq files, we provide a companion script using *Jellyfish* (Marçais and Kingsford, 2011) and *DE-kupl joinCounts* (Audoux et al., 2017) (https://github.com/Transipedia/KaMRaT/tree/master/related-tools).

The *KaMRaT* repository includes different workflow examples and a toy dataset (https://github.com/Transipedia/KaMRaT/tree/master/toyroom/) composed of 89,768 *k*-mers × 20 samples from 10 lung tumor and 10 normal lung samples from a public dataset (Seo et al., 2012) (kmer-counts.subset4toy.tsv.gz), and a (sample, condition) file (sample-condition.toy.tsv).

Typical use cases with these data include:

~~~
# Always Index as first step
kamrat index -intab kmer-counts.subset4toy.tsv.gz -outdir kamrat.idx \
       -klen 31 -unstrand -nfbase 1000000000
# Case 1: merge k-mers into contigs, then select top 5% contigs
#     by t-test p-values between conditions.
kamrat merge -idxdir kamrat.idx -outpath merged-kmers.bin
kamrat score -idxdir kamrat.idx -scoreby ttest.padj -design sample-condition.toy.tsv \
       -seltop 0.05 -with merged-kmers.bin -outpath top-ctg-counts.tsv -withcounts
# Case 2: score k-mers by Spearman correlation to a value vector,
#     then merge the top 5000 k-mers into contigs.
kamrat score -idxdir kamrat.idx -scoreby spearman -design sample-values.toy.tsv \
       -seltop 5000 -outpath top-kmers.bin
kamrat merge -idxdir kamrat.idx -overlap 30-15 -with top-kmers.bin:min \
       -outpath top-ctg-counts.tsv -withcounts mean
# Case 3: fetch k-mers with >= 3 counts in >= 5 “up” samples
#     and zero counts in “down” samples, as labelled in an external list,
#     then merge them into contigs.
kamrat filter -idxdir kamrat.idx -design sample-up.toy.tsv \
       -upmin 3:5 -downmax 0:10 -outpath top-kmers.bin
kamrat merge -idxdir kamrat.idx -overlap 30-15 -with top-kmers.bin:min \
       -outpath top-ctg-counts.tsv -withcounts mean
# Case 4: fetch k-mers absent in a reference (novel k-mers)
     then merge them into contigs.
kamrat mask -idxdir kamrat.idx -fasta gencode.v34.fa -outpath masked-kmers.bin
kamrat merge -idxdir kamrat.idx -overlap 30-15 -with masked-kmers.bin:min \
       -outpath novel-ctg-counts.tsv -withcounts mean
# Case 5: obtain counts for sequences in set, computed as median
#     of component k-mer counts.
kamrat query -idxdir kamrat.idx -fasta sequences.fa -toquery median \
       -outpath query-ctg-counts.tsv
~~~

As an example, the first case reduces the table from 89,768 *k*-mer features to 4,792 contig features, among which 68 have a t-test adjusted p-value *<* 0.05. Using the -seltop option provided by *KaMRaT score*, users can keep for example top 5% contigs (i.e., 240) regardless of their p-value for further investigation.

#### Dependencies

*KaMRaT* is implemented in C++ with dependencies on *MLpack* (Curtin et al., 2018), *Armadillo* (Sanderson and Curtin, 2016) and *Boost Iostreams* (Schäling, 2011) libraries. Source code is available on a GitHub repository https://github.com/Transipedia/KaMRaT.

### Benchmarking

#### Hardware

Benchmarks were performed on a PBS cluster using a single node CPU (Intel Xeon E5-2640v4 @ 2.40GHz, 750Gb memory) with network-mounted hard disk drive (I/O times clocked at 258MB/s).

#### D1 - synthetic error-free RNA-seq reads

This dataset contains 20 samples, synthesized based on Gencode transcripts longer than 5500 nt. We applied *Mutation-Simulator* (Kühl et al., 2021) on this reference to create 10 individual transcriptomes with mutations and applied the *polyester* R package (Frazee et al., 2015), requesting error-free sequence reads with a sequencing depth of 0.2 to emulate discontinuous coverage.

#### D2 - real RNA-seq dataset

This dataset is composed of 154 lung adenocarcinoma RNA-seq samples (77 tumors and 77 adjacent normal tissues) from SRA accession ERP001058 (Seo et al., 2012), processed by *Cutadapt* (Martin, 2011) for adapter and low-quality bases trimming.

#### *k*-mer count matrices

The *k*-mer count matrices for datasets **D1** (149M *k*-mers) and **D2** (484M *k*-mers) were produced using our script https://github.com/Transipedia/KaMRaT/tree/master/related-tools/prepare_kmer_table. For dataset **D2**, subsets corresponding of 5 to 100 samples were produced by selecting 5 to 100 columns in the larger matrix and deleting *k*-mers with zero sum counts.

#### Benchmark of *KaMRaT* for *index, merge* and *score*

Runtime, memory and disk usage were first evaluated on *k*-mer matrices from real dataset **D2** with 5 to 154 samples (Figure S1A). However in this test, both the number of *k*-mers and the number of counts for each *k*-mer increases with the number of samples. To observe the effect of the number of *k*-mers or features alone, we took the **D2** matrix for 50 samples and downsampled it keeping 3.5M to 420M *k*-mers. This allowed us to observe CPU time and RAM usage as a function of the number of *k*-mers or features, at a constant number of samples (Figure S1B).

**Figure S1:**
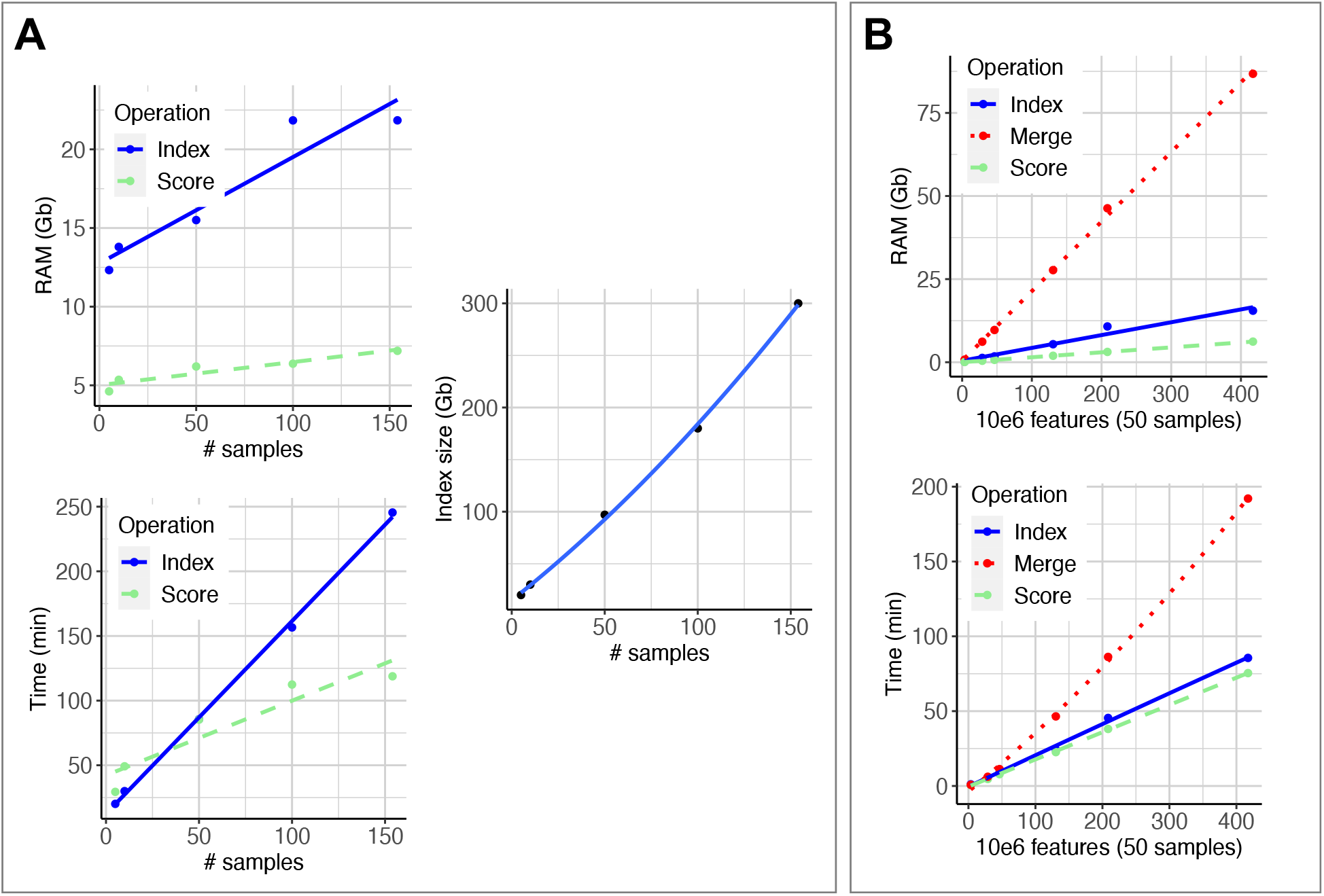
Memory, disk and CPU usage of the main *KaMRaT* modules. A: CPU time, memory and disk usage for *index* and *score* as a function of the number of samples in the **D2** real RNA-seq dataset. B: CPU time and memory usage for *index, score* and *merge* as a function of the number of *k*-mers (features) in the **D2** real RNA-seq dataset.

#### Evaluation of *merge* intervention modes

To assess the correctness of *KaMRaT merge* contigs, we used (i) the synthetic dataset **D1** where all reads were produced from a known set of transcripts and (ii) the real dataset **D2** restricted to 50 samples and further processed by *KaMRaT score* (T-test, “tumor” *vs*. “normal”) keeping the top 1 million *k*-mers scored by T-test P-value. This last test emulated a typical situation where contigs are produced from a condition-specific list of *k*-mers.

*KaMRaT index-merge* was run with different intervention options (none and each intervention method: Mac:0.2, Pearson:0.2, Spearman:0.2). We then compared contigs produced by the different intervention options using Venn diagrams (Figure S2A. Next, we evaluated the correctness of contigs that were specific to each intervention method, by aligning contigs using *BLASTn* (Camacho et al., 2009) to either the transcript set used for simulating reads (simulated dataset **D1**) or to Gencode V34 transcripts (real dataset **D2**). Using *BLASTn* results, we measured the **perfect match ratio** as the percentage of contigs that were completely and exactly aligned - from first to last nucleotide, without any gap or mismatch - to a transcript in the reference (Figure S2B). Note that the lower perfect match ratio in the real dataset is expected as many contigs correspond to transcripts that are not in Gencode or contain polymorphisms or sequencing errors. Finally, Figure S2C shows the average contig lengths for each category of option-specific contigs.

**Figure S2:**
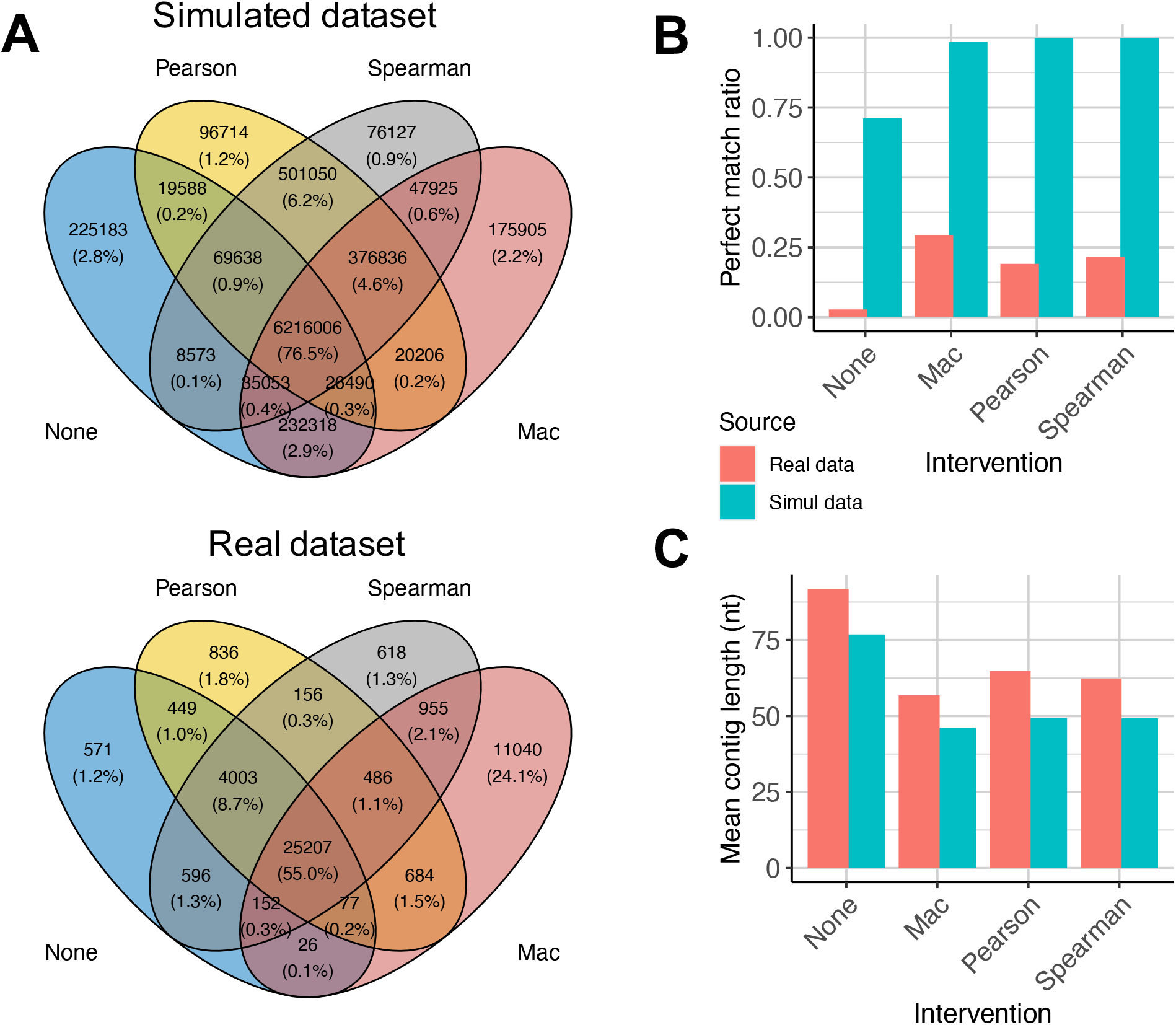
Effect of *merge* intervention modes. A: Venn diagram of contigs produced by the different intervention options, applied to the simulated (**D1**) and real (**D2**, subsampled) datasets. B: Perfect match ratio (contig perfectly aligned to reference over the contig length) of contigs identified specifically by each intervention option. C: Average length of contigs identified specifically by each intervention option.

### Details on scoring methods

The features are scored by different methods in *KaMRaT score* and selected as described below:

**ttest.padj** scores features according to a binary label. Firstly a log_2_(*x* + 1) transformation is applied to sample counts. Then each feature association between sample counts and labels is evaluated by p-value t-test, adjusted by Benjamini-Hochberg procedure for controlling false discovery rate. The top features are selected from the lowest value to the highest.

**ttest.pi** scores features according to a binary label. It is calculated with the formula given in (Xiao et al., 2014) as shown in equation 3. The top features are selected from the highest value to the lowest.

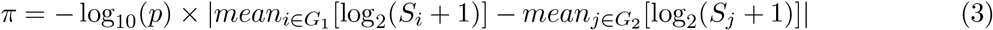

where *p* is the non-adjusted p-value; *G*_1_ and *G*_2_ are two sample groups; *S*_*i*_ and *S*_*j*_ are counts of two samples from *G*_1_ and *G*_2_, respectively.

**snr** scores features according to a binary label. It is calculated by dividing the difference between group means by the sum of group standard deviations, followed by what proposed in (Golub et al., 1999), as shown in equation 4. The top features are selected with absolute values from highest to lowest.

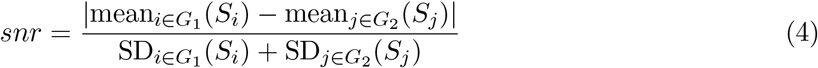

where *G*_1_ and *G*_2_ are two sample groups; *S*_*i*_ and *S*_*j*_ are counts of two samples from *G*_1_ and *G*_2_, respectively.

**dids** scores features according to a multiple-condition label. We generalized the initial case-vs-control version proposed in (de Ronde et al., 2013) to multiple conditions, by iteratively estimating the DIDS score under “one-group-vs-others” fashion and returning the maximum value as the final score, as shown below. The top features are selected from the highest to the lowest value.

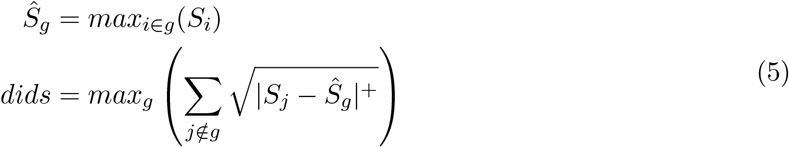

where *g* is a group label; *S*_*i*_ is the count of sample *i*; |*x*|^+^ is *x* if *x >* 0 or 0 otherwise.

**lr** scores features according to a binary label. It estimates classification accuracy by logistic regression, calculated by *MLPack* (Curtin et al., 2018) library. The “lr” scoring method applies a standardization preprocessing of feature counts, i.e., substract all components of the sample count vector with their mean value, and divide them by the standard deviation. The top features are selected from highest to lowest value.

**bayes** scores features according to a multiple-condition label. It estimates classification accuracy using a Bayes classifier from the *MLPack* (Curtin et al., 2018) library. The “bayes” scoring method applies also a standardization preprocessing to feature counts. The top features are selected from highest to lowest score.

**pearson** and **spearman** score features using a vector of continuous values across samples (target numerical vector). These estimate Pearson or Spearman correlation between each feature’s sample count vector and the given target numerical vector, followed by calculation of the absolute value. This option finally selects the top features ranked from highest to lowest score.

**sd** scores features without considering labels (i.e., non-supervised feature selection). It estimates each features’ standard deviation across samples, and selects them from highest to lowest score.

**rsd1** and **rsd2** score features without considering labels (i.e., non-supervised feature selection). They estimate each features’ relative standard deviation across samples to the mean sample count (i.e., SD divided by mean value) or to the minimum sample count (i.e., SD divided by minimum value), respectively. Please note that when the mean or minimum count is less than 1, the original SD is returned without scaling. The methods select top features ranked from highest to lowest score.

**entropy** scores features without considering labels (i.e., non-supervised feature selection). It estimates each features’ Shannon entropy across samples. In order to eliminate the interference of the 0 component in the vector, we added an offset 1 to each component of the sample count vector before calculating the entropy score. This method is used in the *iMOKA reduce* step for informative feature filtering, though *iMOKA* did not add the offset to the count vector, but skipped the 0 components. (Lorenzi et al., 2020). The top features are selected from smallest to largest score.

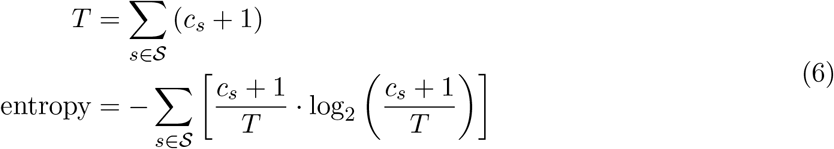

where 𝒮 is the sample set, *c*_*s*_ is the count of sample *s*.

#### Comparison of scoring methods

To compare *score* methods, we generated four feature count matrices for 300 samples and 20,000 to 200,000 features, using the *compcodeR* R package (Soneson, 2014). The four matrices differ from each other by the level of differential expression (DE) signal intensities and noises, as indicated in Figure S3A. Using the simulated DE status of features as ground truth, we evaluated the DE detection capability of each scoring method applicable to categorical variables, i.e., t-test adjusted p-value and *π*-value, SNR, DIDS, logistic regression and Bayes classifier, using precision-recall curves (Figure S3B). DIDS predictions were most distant from other scoring methods (Figure S3C). T-test adjusted p-values and *π*-values achieved the best performance, with respective PR AUCs of 0.794 and 0.785 in the most complicated case. SNR also showed favorable performance with a worst-case PR AUC of 0.637. These three methods estimate the difference between groups based on means and standard deviations, which coincides with how *compcodeR* simulates DE features. While other methods (Bayes, DIDS and LR) did not perform as well, it should be noted that Bayes and LR are not intended to detect DE features. In principle DIDS detects DE features by considering outlier observations, however the way outliers are generated in *compcodeR* is independent from differential feature labeling, thus it is of no use for DIDS, which probably explains its behavior.

**Figure S3:**
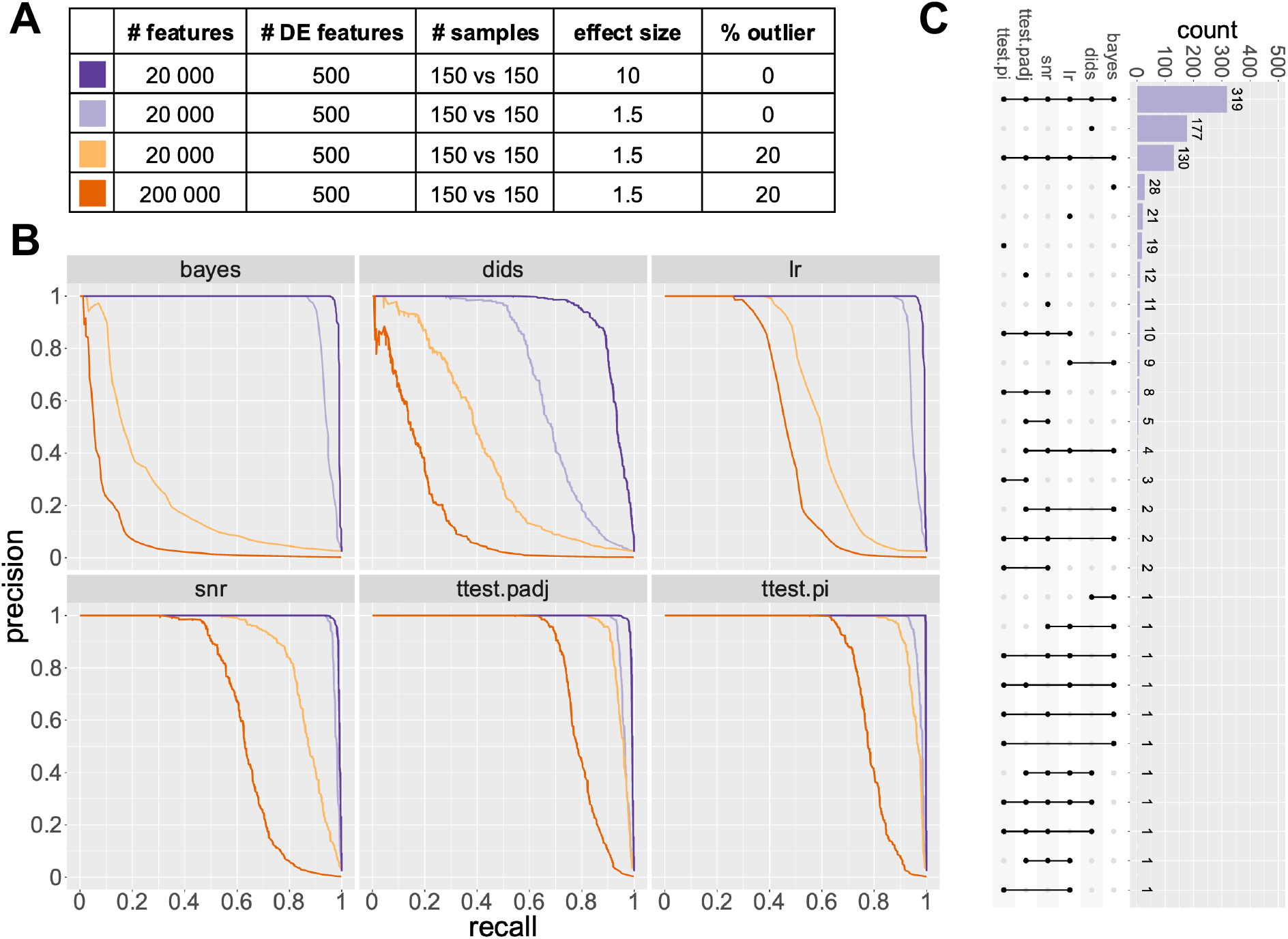
Comparison of *score* methods (color codes are defined in panel A). A: Characteristics of simulated count matrices. B. Precision-recall curves for classification of DE features by each scoring method. C. UpSetR plot comparing predicted DE features from each method, for the second simulated dataset (20,000 features, effect size 1.5, no outlier).

